# TifBERT: a self-supervised foundation model for normalization-robust bulk RNA-seq representation learning

**DOI:** 10.64898/2026.06.08.728683

**Authors:** SeyedMohsen Hosseini, Divya Sharma

## Abstract

Bulk RNA sequencing remains central to translational genomics, yet foundation-model development has largely focused on single-cell data. Existing transformer approaches for bulk RNA-seq often rely on expression discretization, numerical reconstruction, external gene embeddings, or restricted gene sets, limiting robustness across normalization schemes and cohorts. Here, we introduce TifBERT, a self-supervised framework for full-transcriptome bulk RNA-seq representation learning. TifBERT converts each unordered expression profile into a sample-specific gene sequence using term frequency-inverse document frequency (TF-IDF) ordering, prioritizing genes that are both highly expressed within a sample and selectively expressed across the cohort. It is then pretrained using masked gene modeling, predicting gene identities from transcriptomic context rather than reconstructing expression values.

Pretrained on harmonized TCGA Pan-Cancer data spanning five RNA-seq normalization schemes, TifBERT learns contextual representations across approximately 10,000 genes without expression binning, landmark-gene restriction, or external biological embeddings. Across 33 TCGA cancer types, TifBERT achieved 90.83% accuracy, 0.996 macro AUC-ROC, and 0.903 MCC. It also captured pathway-level biology, achieving mean sample-wise and pathway-wise Pearson correlations of 0.754 and 0.762 across 1,387 PARADIGM pathway activities. Independent evaluation on GTEx healthy tissues showed preservation of tissue-level transcriptomic structure without retraining. In comparison with existing models, TifBERT achieves competitive subtype discrimination with substantially greater stability and produces markedly richer embedding geometry (effective rank 95.6 versus 6.3), without requiring expression discretization or in-distribution pretraining exposure. Together, TifBERT provides a scalable, normalization-independent foundation model for reusable bulk transcriptomic representation learning.

## Introduction

Bulk RNA sequencing (RNA-seq) remains one of the most widely used molecular profiling technologies in translational genomics, clinical cohort studies, and public repositories. It has enabled disease classification, patient stratification, pathway analysis, prognosis modeling, and discovery of tissue- and disease-specific transcriptional programs^1–6^. Yet, despite the rapid emergence of foundation models for biological data, bulk RNA-seq still lacks a broadly reusable self-supervised representation model that can operate at transcriptome scale, transfer across cohorts and normalization schemes, and support diverse downstream tasks without extensive task-specific feature engineering.

Modeling bulk RNA-seq is challenging because each sample is represented by tens of thousands of gene-expression measurements, whereas most studies contain far fewer samples than features. This high-dimensional, low-sample-size regime makes supervised deep learning prone to overfitting and has motivated the use of dimensionality reduction, predefined gene panels, pathway summaries, or manually selected features. Although these strategies improve tractability, they can discard weak, context-dependent, or disease-specific signals distributed across the transcriptome. Generative models, including variational autoencoders, generative adversarial networks, and diffusion models, have been used to learn transcriptomic latent spaces and generate synthetic expression profiles^7–10^. However, many such approaches remain sensitive to preprocessing choices, require careful stabilization, or struggle to scale to whole-transcriptome modeling in a way that yields reusable contextual representations.

Transformers^11^ offer an alternative paradigm, where instead of reducing gene-expression profiles to fixed feature vectors or predefined gene sets, they can learn contextual relationships among genes through self-supervised pretraining. This strategy has transformed representation learning in natural language and has increasingly been applied to biomedical, molecular, and single-cell data^12–15^. However, transcriptomes pose a fundamental tokenization problem. Unlike words in a sentence or residues in a protein, genes in a bulk RNA-seq profile do not have a canonical sequence order. A transformer for bulk transcriptomics must therefore define an ordering strategy that is biologically informative, robust to normalization, and not overly dependent on raw expression magnitudes.

Single-cell transcriptomic foundation models have made substantial progress toward this goal. Geneformer uses rank-based gene ordering to support transfer learning across network biology tasks^16^, scGPT learns generative representations from millions of single-cell profiles^17^, and GenePT constructs gene and cell embeddings using language-model representations of gene descriptions^18^. These methods show that self-supervised pretraining can produce transferable representations of transcriptomic state. However, they are designed for single-cell data, where each profile reflects an individual cellular state and where cell-to-cell heterogeneity provides a rich source of variation. Bulk RNA-seq aggregates expression across many cells and tissue compartments, producing a different statistical object from single-cell profiles. It exhibits distinct normalization dependencies, lower within-cohort expression granularity, and is frequently derived from retrospective clinical cohorts. As a result, single-cell tokenization and pretraining strategies cannot be directly assumed to solve the representation-learning problem for bulk RNA-seq.

Recent efforts have begun to adapt transformer pretraining to bulk transcriptomics. BulkRN-ABert applies masked language modeling to TCGA bulk RNA-seq after discretizing continuous expression values and incorporating Gene2Vec embeddings^19,20^. GexBERT represents each gene using both gene identity and measured expression and trains the model to reconstruct held-out expression values^21^. These approaches demonstrate the promise of transformer-based bulk RNA-seq modeling, but they also expose a central limitation that representation learning remains closely coupled to numerical expression magnitudes, either through discretized expression tokens or expression-reconstruction objectives. This coupling reduces portability across datasets generated with different preprocessing pipelines, normalization schemes, and measurement scales. In parallel, landmark-gene approaches such as LINCS L1000 improve efficiency by restricting the input space, but at the cost of excluding much of the transcriptome, including low- and moderately expressed genes that may carry regulatory or disease-specific information^22,23^.

Here, we introduce TifBERT, a self-supervised transformer framework for full-transcriptome bulk RNA-seq representation learning. TifBERT is built around three methodological principles. First, it converts each unordered expression profile into a sample-specific gene sequence using term frequency-inverse document frequency (TF-IDF) ordering. This ordering prioritizes genes that are both abundant within a sample and selectively expressed across the cohort, allowing the model to emphasize sample-specific transcriptional programs while retaining transcriptome-wide coverage. Second, TifBERT pretrains the encoder using masked gene modeling, where masked gene identities are predicted from their ordered transcriptomic context, rather than by reconstructing numerical expression values. This decouples representation learning from expression scale and reduces dependence on dataset-specific normalization. Third, TifBERT operates across approximately 10,000 genes using overlapping sliding windows, enabling whole-transcriptome contextual learning without predefined landmark genes, external gene embeddings, or projection-based compression.

Together, these design choices position TifBERT as a reusable foundation-model framework for bulk transcriptomics, addressing a key methodological gap between the rapid progress of single-cell foundation models and the continued centrality of bulk RNA-seq in clinical and translational genomics.

## Methods

TifBERT is designed as a self-supervised representation learning framework for bulk RNA-seq, addressing the challenge of converting unordered, high-dimensional expression profiles into transferable contextual representations without dependence on expression-value reconstruction or dataset-specific normalization. The framework consists of four components: TF-IDF-based gene ordering, sliding-window sequence construction, masked gene modeling pretraining, and attention-based sample-level aggregation. An overview of the complete framework is presented in Figure 1, and the following subsections describe each component in detail.

**Figure 1.**
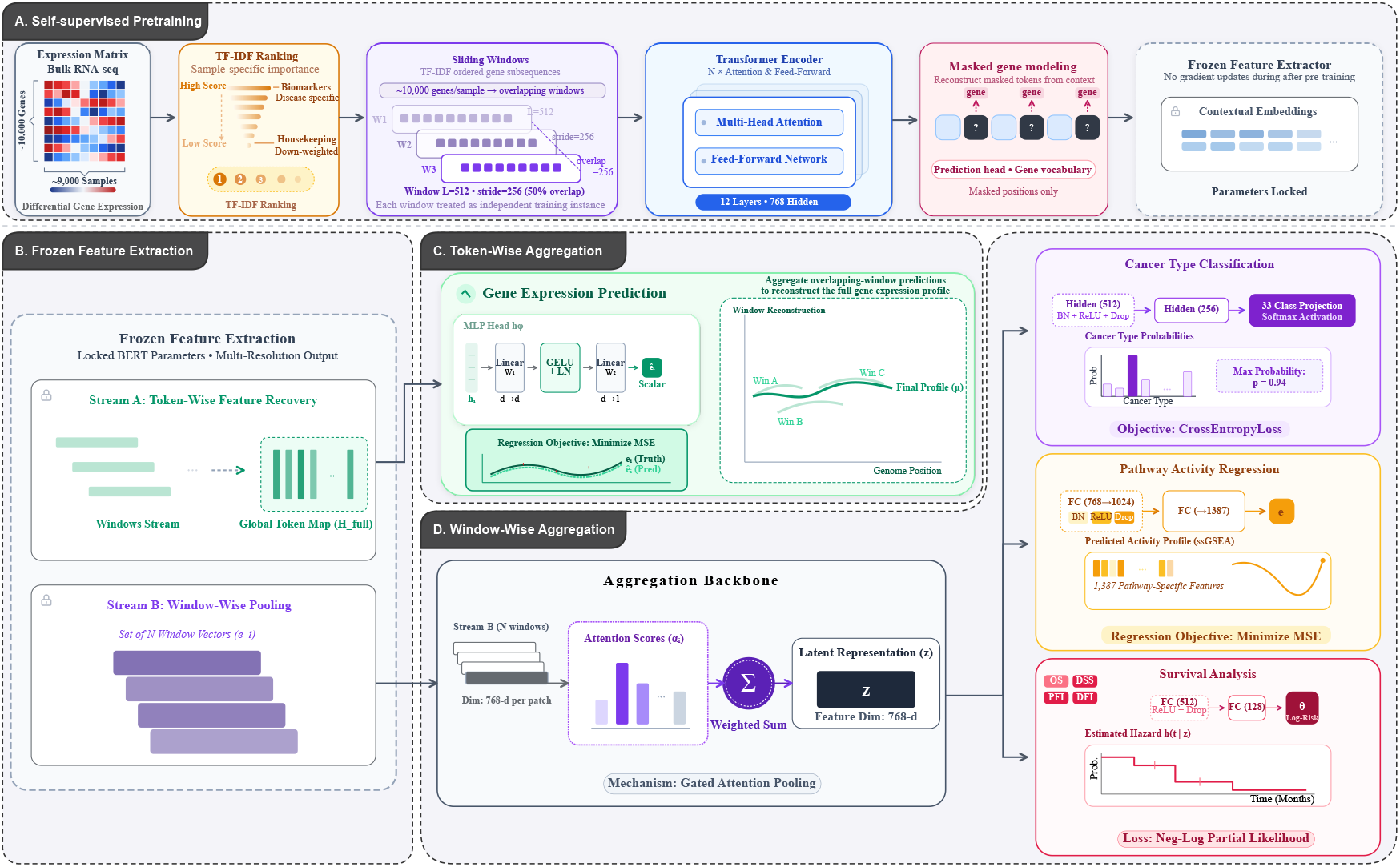
Overview of the TifBERT framework. (A) Self-supervised pretraining using a masked gene modeling objective on TF-IDF-ordered bulk RNA-seq sequences, where randomly selected gene tokens are masked and reconstructed from contextual transcriptomic information. (B) Frozen feature extraction, where the pretrained transformer encoder is kept fixed and used to compute contextualized gene embeddings for each sliding window without task-specific fine-tuning. (C) Token-wise aggregation, where token-level hidden states within each window are aggregated via mean pooling to obtain a fixed-length window representation that summarizes local transcriptomic context. (D) Window-wise aggregation, where overlapping window embeddings are combined into a single sample-level representation, enabling robust downstream prediction across classification, regression, and survival analysis tasks.

### Study design and data sources

TifBERT was developed as a self-supervised representation learning framework for bulk RNA-seq profiles. The model was pretrained using harmonized transcriptomic profiles from the TCGA Pan-Cancer Atlas and the UCSC Toil RNA-seq Recompute resource^24,25^. To encourage robustness to preprocessing variation, five RNA-seq representations were included: batch-effect normalized TCGA expression profiles and four Toil-derived quantifications, expected counts, FPKM, TPM, and normalized counts. All samples were mapped to a common gene space before model training.

Gene identifiers were converted from Ensembl IDs to HUGO gene symbols. Genes were retained only if they were present across all five RNA-seq representations. Low-information genes were removed using median-expression and variance-based filters, yielding a shared transcriptomic feature space of approximately 10,000 genes. Full filtering criteria and dataset summaries are provided in Supplementary S1.

The TCGA cohort comprised 9,201 samples in total and was split at the sample level into training, validation, and held-out test sets using a 90%/ 5%/ 5% partition, corresponding to 8,315 training samples, 406 validation samples, and 480 test samples. The same split was used for pretraining and downstream evaluation to prevent leakage across tasks. TF-IDF statistics were computed within each split, ensuring that test-sample gene orderings were derived independently from training data.

### TF-IDF gene ordering

Bulk RNA-seq profiles are unordered vectors, whereas transformer architectures require an ordered sequence of input tokens. TifBERT addresses this by converting each sample into a sample-specific gene sequence using term frequency–inverse document frequency (TF-IDF) ordering. For gene *g*_*i*_ in sample *s*, term frequency was defined as the relative abundance of the gene within that sample,

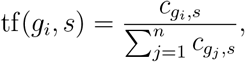

where *c*_*gi,s*_ denotes the expression value of gene *g*_*i*_ in sample *s*. The inverse document frequency term quantified the cohort-level specificity of each gene,

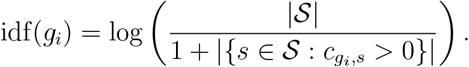

The final score was given by

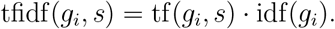

Genes were ranked in decreasing order of their TF-IDF scores to produce a sample-specific sequence. This ordering prioritizes genes that are highly represented within a sample but selectively expressed across the cohort, while retaining transcriptome-wide coverage. Unlike expression-value ranking, TF-IDF ordering incorporates both within-sample abundance and across-sample specificity. Unlike landmark-gene approaches, it does not discard genes before model input.

### Sequence construction

After TF-IDF ordering, each sample was represented as a ranked sequence of gene identities. Because the full transcriptome exceeds the input length of standard BERT-style encoders^26^, each ordered gene sequence was divided into overlapping windows. The primary model used windows of length *L* = 512 with a stride of 256 genes. This overlap allowed genes near window boundaries to be observed in alternative local contexts and reduced edge effects during representation learning and downstream prediction.

Each window was treated as a training instance during pretraining. During downstream evaluation, window-level embeddings were aggregated to form a sample-level representation.

### Self-supervised masked gene modeling

TifBERT was pretrained using a masked gene modeling objective. For each input window, 15% of gene-token positions were selected at random and replaced according to the standard BERT masking strategy: 80% of selected positions were replaced with a special `[MASK]` token, 10% with a randomly sampled gene token, and the remainder left unchanged. The transformer encoder was trained to predict the masked gene identities from the remaining TF-IDF-ordered transcriptomic context:

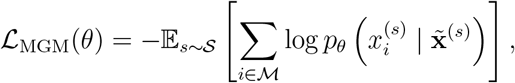

where *ℳ* denotes the set of masked positions, 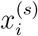 is the original gene token, and 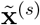 is the masked input sequence.

This objective trains the encoder to learn contextual gene relationships without reconstructing numerical expression values during pretraining. By predicting gene identities, this objective decouples the pretrained representation from expression scale, discretization scheme, and normalization pipeline.

### Transformer encoder

The TifBERT encoder uses a BERT-style architecture^26^ with 12 transformer layers, hidden dimension 768, 12 attention heads, and intermediate feed-forward dimension 3072, totaling approximately 93M parameters. The model was pretrained on TF-IDF-ordered gene windows using AdamW with a 1*e* − 4 learning rate schedule for 350 epochs on 4 NVIDIA H100 GPUs, requiring approximately three days. The encoder was subsequently frozen for all downstream tasks, separating representation learning from task-specific optimization and ensuring that downstream performance reflects the quality of pretrained representations rather than task-specific fine-tuning. This design also reduces the risk of overfitting in task-specific supervised training given the relatively small sample sizes typical of clinical transcriptomic cohorts. Alternative model capacities, sequence lengths, ordering strategies, and aggregation schemes were evaluated during development; full ablation results and training configuration details are provided in Supplementary S2.

### Sample-level representation

For downstream analyses, each sample was represented by aggregating frozen encoder outputs across its sliding windows. Each window was processed independently, and window-level representations were obtained by mean-pooling token embeddings within each window. These window embeddings were then combined into a fixed-dimensional sample embedding, **z**^(*s*)^ *∈* R^768^, using a gated attention pooling module, which computed a weighted sum of window embeddings with attention weights derived from a learned scoring function applied to each window representation. This shared embedding was used across all downstream tasks including classification, pathway regression, expression prediction, and survival analysis. Details of alternative aggregation strategies are provided in Supplementary S2.

### Downstream evaluation

The pretrained encoder was evaluated using task-specific heads trained on top of frozen TifBERT representations, with all encoder parameters held fixed during downstream training. Full architecture specifications, loss functions, and training configurations for each task are provided in Supplementary S2.

For cancer-type classification, sample embeddings were passed to a multilayer perceptron trained to predict 33 TCGA cancer types using cross-entropy loss. For pathway activity regression, sample embeddings were used to predict 1,387 PARADIGM pathway activity scores obtained from the UCSC Xena Pan-Cancer Atlas Hub, corresponding to z-score-transformed ssGSEA pathway activities integrating pathway information from NCI-PID, BioCarta, and Reactome^27–30^, optimized using mean squared error loss. For gene-expression prediction, token-level encoder outputs were passed through a lightweight regression head to estimate expression values per gene within each window; when a gene appeared in multiple overlapping windows, predictions were averaged across all windows containing that gene. For survival prediction, separate Cox-style neural models were trained for overall survival, disease-specific survival, progression-free interval, and disease-free interval, each producing a scalar log-risk score optimized via negative log partial likelihood. Full endpoint definitions, censoring details, and task-specific architectures are provided in Supplementary S2.

### External GTEx Validation

To assess transfer beyond the TCGA cancer distribution, the pretrained TifBERT encoder was evaluated on an independent subset of 500 GTEx healthy tissue samples spanning 27 tissue types, selected as described in Supplementary Table S2. No GTEx samples were used during pretraining, validation, model selection, or hyperparameter tuning. GTEx data were processed using the same gene harmonization pipeline as TCGA and Toil data, with expression values transformed using log_2_(count + 1) to match the expected-count representation. Tissue-level transcriptomic structure was evaluated by computing sample embeddings and applying Procrustes alignment to compare cancer-type centroid configurations derived from actual and predicted expression profiles, as described in Supplementary S2. This evaluation assesses structural preservation of tissue-level organization rather than task-specific predictive performance on GTEx samples, which remains an avenue for future work.

### Comparative evaluation

TifBERT representations were compared against BulkRNABert^19^ and a non-contextual top-K TF-IDF baseline across cancer-type classification, embedding geometry, and neighborhood retrieval metrics. For BulkRNABert, publicly available pretrained weights were used. Full evaluation protocols, metric definitions, and implementation details for all comparisons are provided in Supplementary S2.

## Results

TifBERT was evaluated as a frozen self-supervised encoder across five tasks: cancer-type classification, gene-expression prediction, pathway activity regression, survival prediction, and external transfer to GTEx healthy tissues. Unless otherwise stated, all results are reported on held-out test samples not used during pretraining, validation, or model selection.

### Cancer-type classification

TifBERT achieved high discrimination across 33 TCGA cancer types using frozen encoder embeddings and a lightweight classification head. The best-performing configuration used mean-pooled window embeddings with attention-based aggregation, reaching 90.83% accuracy, a macro AUC-ROC of 0.9965, and an MCC of 0.9036 on the held-out test set (Table 1). The correct cancer type appeared among the top three predictions for 97.29% of samples and among the top five for 98.96% of samples. Full per-class F1-score analysis and ROC curves are provided in Supplementary S3.

**Table 1.**
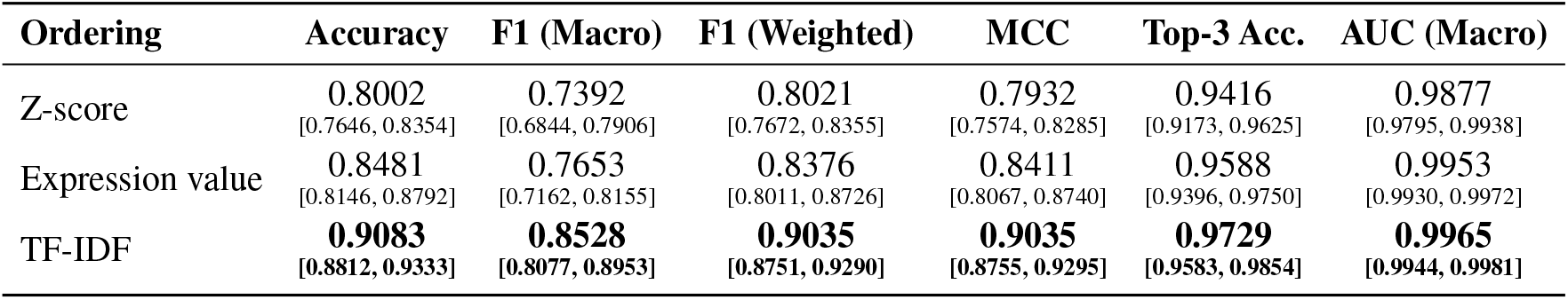
Cancer-type classification performance across gene ordering strategies using the base TifBERT architecture and mean-attention aggregation. Results are reported with 95% bootstrap confidence intervals on the held-out test set.

To assess whether performance depended on the proposed ordering strategy, TF-IDF ordering was compared with expression-value and Z-score ordering using an identical encoder and classification head. TF-IDF ordering achieved the highest performance across all metrics, improving accuracy from 84.81% with expression-value ordering and 80.02% with Z-score ordering (Table 1). The consistent advantage of TF-IDF indicates that cohort-level gene specificity provides an informative ordering signal beyond expression magnitude alone.

### Comparison with baselines

To determine whether performance gains arise from contextual representation learning rather than gene identity alone, TifBERT was compared against a non-contextual top-K gene-set baseline. In this baseline, the 200 highest-ranked TF-IDF genes per sample, were encoded as a binary indicator vector and used to train a logistic regression classifier. The top-200 logistic regression baseline achieved 65.00% accuracy and an AUC-ROC of 0.9803 (Table 2). TifBERT substantially outperformed this baseline, reaching 90.83% accuracy and an AUC-ROC of 0.9965. This gap demonstrates that masked gene modeling captures gene-to-gene contextual relationships that improve multiclass discrimination substantially beyond what gene identity information alone can provide.

**Table 2.** Comparison of TifBERT with non-contextual baseline for 33-class cancer-type classification on the held-out TCGA test set.

| Model | Accuracy | F1 (Macro) | MCC | AUC-ROC |
| --- | --- | --- | --- | --- |
| Top-200 + LR | 0.6500 | 0.6706 | 0.6475 | 0.9803 |
| TifBERT | **0.9083** | **0.8528** | **0.9035** | **0.9965** |

### Gene expression prediction

Although TifBERT is not pretrained to reconstruct numerical expression values, frozen contextual representations support accurate gene-expression prediction through a lightweight downstream head. Prediction error varies systematically with token position within each sliding window, with higher error near window boundaries and lower error near the window center (Figure 2), consistent with reduced bidirectional context at edge positions. Because adjacent windows overlap by 256 genes, most genes appear in multiple local contexts, and final gene-level predictions are averaged across all windows containing that gene to mitigate boundary effects.

**Figure 2.**
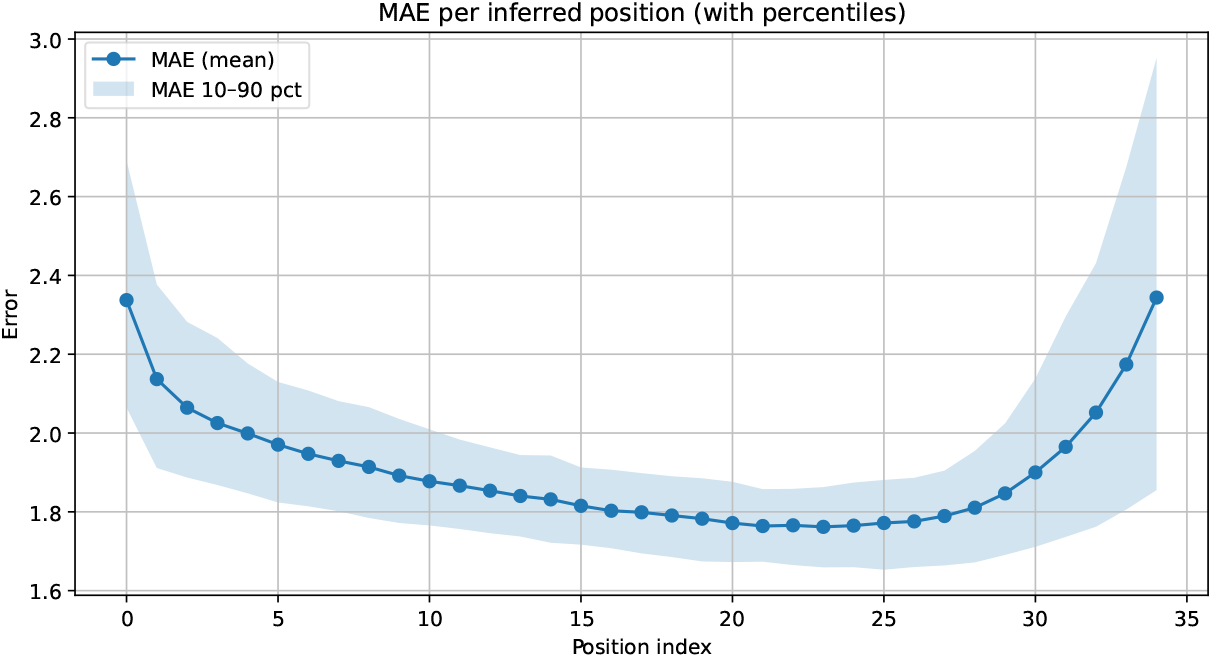
Mean absolute error as a function of token position within a sliding window, averaged across held-out TCGA test samples. Error is highest near window boundaries and lowest near the window center, consistent with reduced bidirectional context at edge positions. Shaded area indicates the 10th–90th percentile range across samples. This positional effect does not propagate to final gene-level predictions because the 256-gene window overlap ensures most genes appear near the center of at least one window.

To assess whether predicted profiles retained higher-order biological structure, actual and predicted gene-expression profiles were jointly embedded using UMAP and annotated by cancer type. Predicted profiles preserved disease-specific organization in the shared manifold (Figure 3), and Procrustes alignment of cancer-type centroids confirmed that predicted expression profiles retained cancer-type-level structure in gene-expression space (Figure 4). Together these results indicate that TifBERT representations encode transcriptomic organization at both the individual gene and disease-program level, despite the absence of expression-value reconstruction during pretraining.

**Figure 3.**
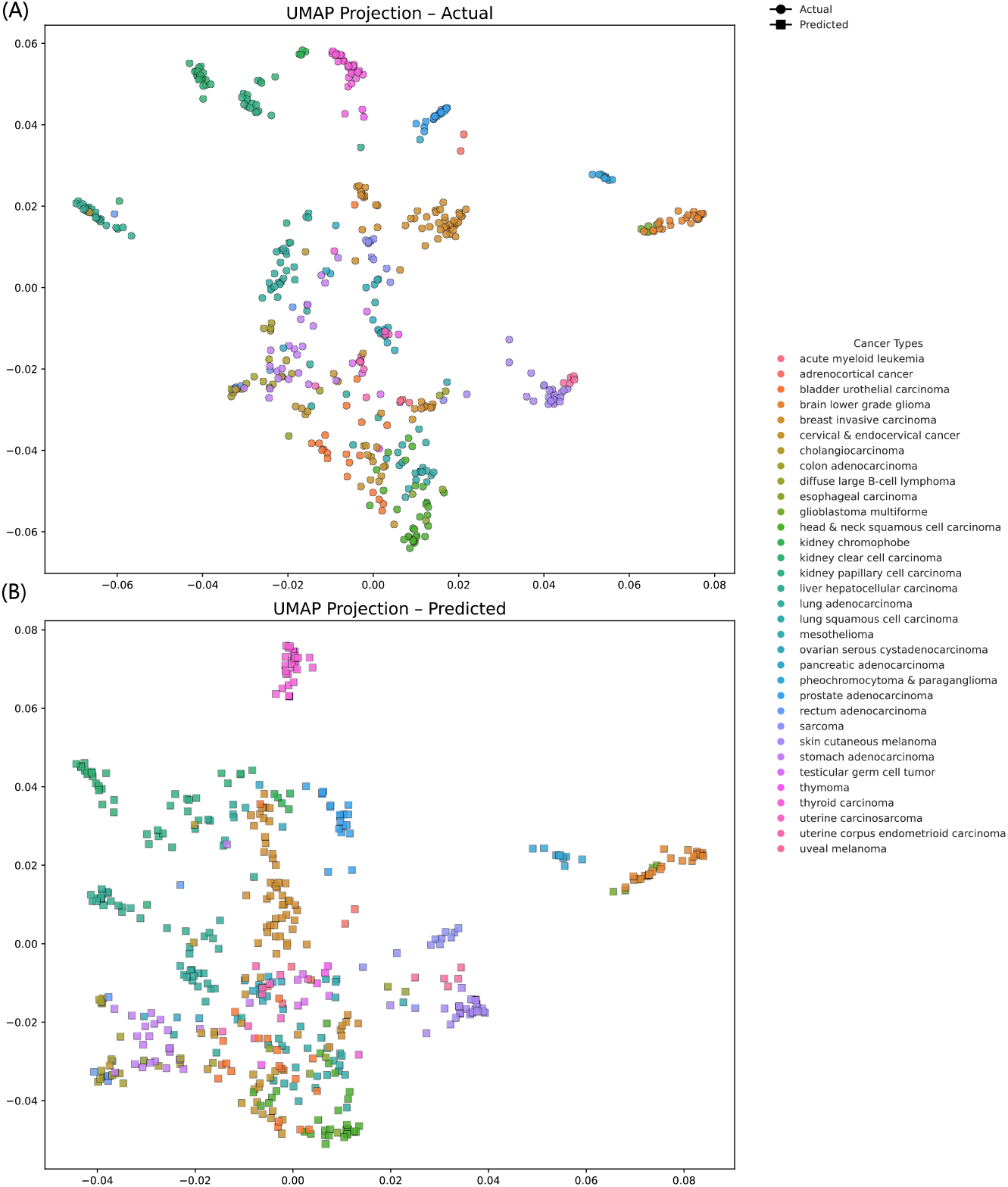
UMAP visualization of gene-expression representations for the held-out TCGA test set (*n* = 480 samples). Actual (A) and model-predicted (B) expression profiles were jointly embedded into a shared low-dimensional manifold using identical preprocessing and UMAP parameters, and samples were colored according to cancer type. Despite predictions being generated from frozen pretrained representations without explicit expression-value reconstruction during pretraining, the predicted profiles recapitulate the global transcriptomic organization observed in real data. Cancer-type clusters remain well separated and preserve relative neighborhood relationships across the manifold, indicating that the model maintains higher-order biological structure rather than merely matching marginal expression statistics.

**Figure 4.**
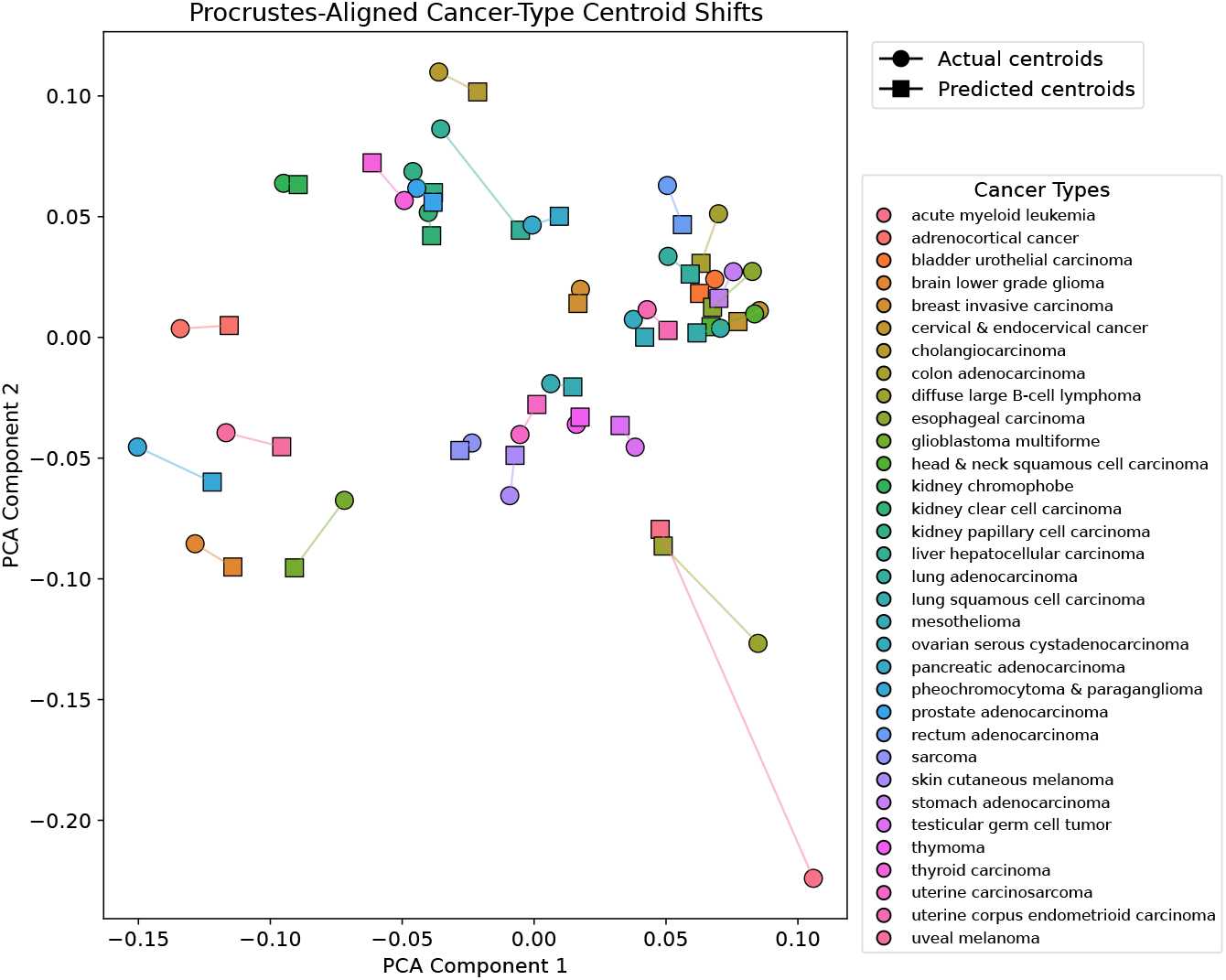
Procrustes alignment of cancer-type centroids derived from actual and predicted gene-expression profiles for the held-out TCGA test cohort (*n* = 480 samples). For each cancer type, sample-level expression profiles were first embedded into a common representation space, and centroid positions were computed separately for actual and predicted data. The two centroid configurations were then aligned using Procrustes analysis to remove differences due to translation, rotation, and scaling. Visualization in the first two principal components shows paired markers corresponding to each cancer type, with connecting segments linking actual and predicted centroids. Short connecting distances across most cancer types indicate strong geometric agreement between the two configurations, demonstrating that predicted expression profiles preserve global disease-specific transcriptomic organization and inter-cancer relationships in gene-expression space.

### Pathway activity regression

TifBERT embeddings were used to predict 1,387 PARADIGM pathway activity scores from frozen sample representations. With TF-IDF ordering, the model achieved a mean sample-wise Pearson correlation of 0.754 and a mean pathway-wise Pearson correlation of 0.762 on the held-out test set (Table 3). The mean sample MAE of 0.494 on z-scored targets corresponds to approximately half a standard deviation of prediction error, which is low given that z-scored pathway scores span a range of approximately [−3, +3]. The near-identical performance across sample-wise and pathway-wise metrics indicates that the learned representations capture pathway co-regulation structure consistently across both samples and biological programs rather than fitting one axis at the expense of the other.

**Table 3.** Pathway activity regression performance on the held-out TCGA test set (480 samples) across 1,387 PARADIGM pathway targets and three gene ordering strategies. Sample-wise Pearson and MAE are averaged across samples; pathway-wise Pearson and MAE are averaged across pathways. TF-IDF ordering achieves the strongest performance on all metrics.

| Ordering | Sample<br>Pearson | Sample<br>MAE | Pathway<br>Pearson | Pathway<br>MAE |
| --- | --- | --- | --- | --- |
| Z-score | 0.7131 | 0.5352 | 0.7300 | 0.5351 |
| Expression value | 0.7161 | 0.5312 | 0.7444 | 0.5310 |
| TF-IDF | **0.7542** | **0.4943** | **0.7621** | **0.4942** |

TF-IDF ordering outperformed expression-value ordering (sample-wise Pearson 0.716, pathway-wise 0.744) and Z-score ordering (0.713 and 0.730, respectively) across all metrics. The consistent ranking of ordering strategies across cancer classification and pathway regression provides evidence that TF-IDF ordering encodes a general-purpose biological signal rather than a task-specific advantage.

### Transfer to healthy tissues

To assess cross-dataset transferability, the pretrained TifBERT encoder was evaluated on an independent subset of 500 GTEx healthy tissue samples spanning 27 tissue types, with no model retraining or parameter updates. Tissue-level transcriptomic structure was evaluated by comparing cancer-type centroid configurations derived from actual and predicted expression profiles using Procrustes alignment (Figure 5). Centroid positions were well preserved across the majority of tissue types, with a mean Procrustes disparity of 0.2065, indicating that TifBERT representations maintain global tissue-level organization under a substantially shifted biological and experimental distribution. 5 tissue types showed larger centroid displacement, potentially reflecting the domain shift between cancer-derived and healty tissue expression profiles. These results demonstrate structural transferability of TifBERT representations to healthy tissue data, though task-specific benchmarking on GTEx samples remains an avenue for future work.

**Figure 5.**
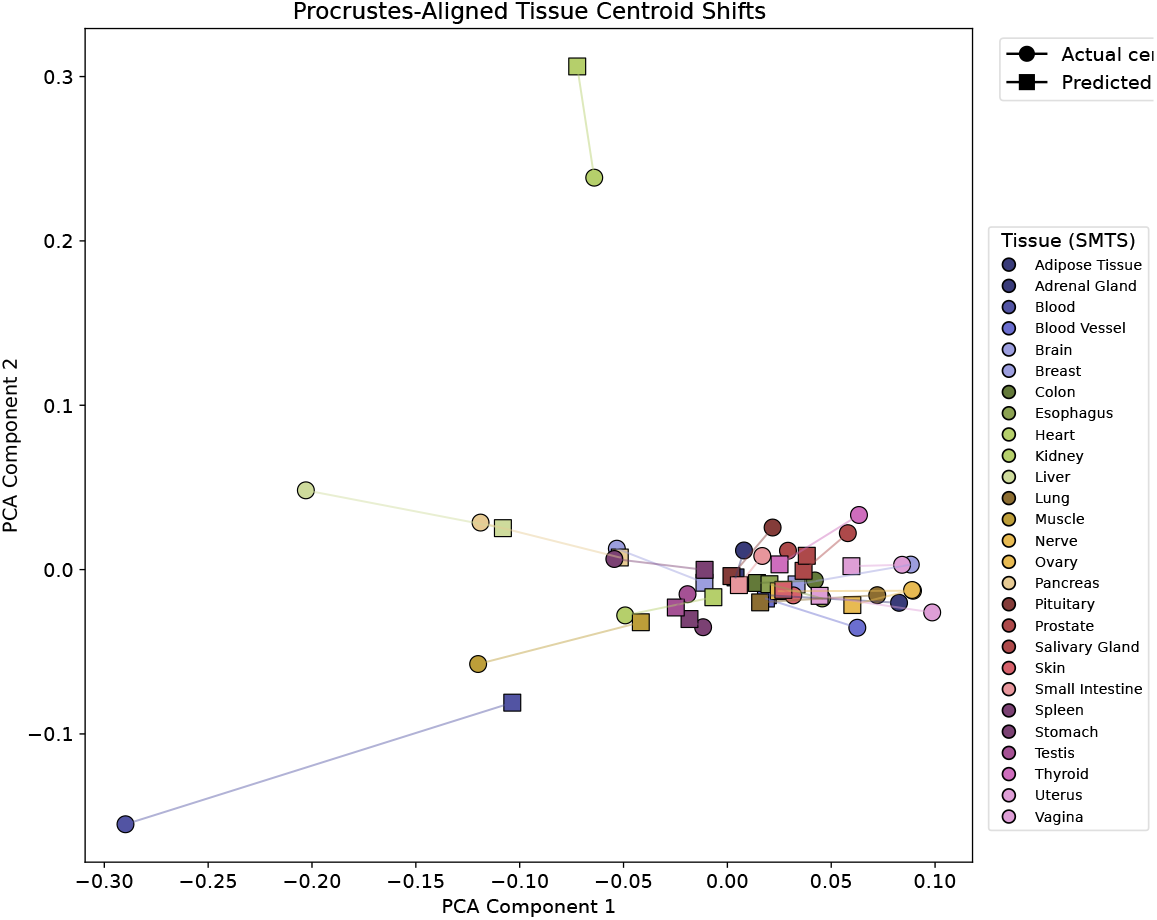
Procrustes-aligned tissue centroids derived from actual and predicted gene-expression profiles for an independent GTEx test set (*n* = 500 samples), visualized in the first two principal components. GTEx samples were not used during pretraining, validation, or model selection, and represent a distinct out-of-distribution setting relative to TCGA. For each tissue type, centroids were computed separately for actual and predicted expression profiles, and paired using Procrustes alignment to remove global geometric transformations. Each connected pair of markers corresponds to a single tissue type, and short connecting distances indicate strong agreement between actual and predicted centroid locations. The observed alignment suggests that TifBERT preserves global tissue-specific transcriptomic structure even under a substantially shifted biological and experimental distribution.

### Survival prediction

TifBERT embeddings were evaluated for time-to-event prediction across four TCGA survival endpoints. With TF-IDF ordering, the model achieved C-index values of 0.646, 0.672, 0.612, and 0.671 for OS, DSS, PFI, and DFI respectively. TF-IDF ordering achieved the highest C-index and mean time-dependent AUC for DSS, PFI, and DFI, while expression-value ordering performed best for OS (C-index 0.684). Z-score ordering consistently underperformed across all four endpoints. Full survival metrics including time-dependent AUC at 1, 2, 3, and 5 years are reported in Supplementary Table S6.

### Summary of ordering strategy effects

Across all evaluated tasks, TF-IDF ordering consistently outperformed expression-value and Z-score ordering for cancer-type classification, pathway activity regression, and three of four survival endpoints. The consistent ranking of ordering strategies across tasks with fundamentally different objectives (multiclass classification, continuous pathway regression, and time-to-event prediction) supports the interpretation that TF-IDF-based gene prioritization captures a general-purpose biological signal rather than a task-specific advantage. These results collectively demonstrate that the combination of sample-specific gene ordering and masked gene identity prediction produces transferable whole-transcriptome representations that support diverse downstream analyses from a single frozen pretrained encoder.

### Representation Quality Evaluation

Representation quality was evaluated on 480 samples spanning 11 cancer types. This dataset is held out from TifBERT pretraining but falls within BulkRNABert’s pretraining distribution, as TCGA data were included during its training. To enable reliable stratified evaluation, only cancer types with at least 20 samples were retained, covering lung, breast, liver, kidney, brain, stomach, thyroid, skin, bladder, and cervix cancers. Representation quality was examined through linear separability, neighborhood retrieval consistency, histological subtype discrimination, and intrinsic embedding geometry (Table 4).

**Table 4.** Representation quality comparison between TifBERT and BulkRNABert on 480 samples across 11 cancer types. Lung subtype accuracy is computed on the 49 lung samples only (adenocarcinoma vs. squamous cell carcinoma). ↑indicates higher is better and ↓ lower is better. ^*†*^Evaluation samples overlap with BulkRNABert’s pretraining data and therefore reflect in-distribution performance.

| Metric | TifBERT | BulkRNABert$^\dagger$ |
| --- | --- | --- |
| *Embedding Geometry* |  |  |
| Effective Rank ↑ | 95.6 | 6.3 |
| Top Eigenvalue Dominance ↓ | 0.114 | 0.585 |
| *Lung Subtype Discrimination (binary probe)* |  |  |
| Accuracy ↑ | $69.3 \pm 6.5\%$ | $71.6 \pm 19.2\%$ |
| *Cross-Cancer Retrieval (Precision@k)* |  |  |
| P@1 ↑ | 0.644 | 0.684 |
| P@5 ↑ | 0.524 | 0.589 |
| P@10 ↑ | 0.460 | 0.519 |
| P@20 ↑ | 0.371 | 0.431 |
| *Linear Probe (5-fold CV)* |  |  |
| Disease Accuracy ↑ | $73.4 \pm 3.8\%$ | $83.4 \pm 3.5\%$ |

### Embedding Geometry

Intrinsic geometry differentiates the models clearly. BulkRNABert showed low effective rank and strong top-eigenvalue dominance, indicating concentration along a small number of dominant directions associated with magnitude reconstruction. TifBERT produced a higher effective rank and lower dominance, corresponding to a more isotropic representation space. These properties reflect differences in pretraining objectives: expression reconstruction promotes axis-aligned compression, whereas masked gene identity prediction over ranked sequences encourages distributed encoding of co-regulatory relationships. Unlike task metrics, these geometric properties are independent of data overlap.

### Linear Probing

A linear classifier was trained on frozen embeddings using 5-fold cross-validation to measure linearly accessible biological information. BulkRNABert achieved higher disease classification accuracy than TifBERT. Two factors contextualize this gap. First, BulkRNABert is pretrained to reconstruct expression magnitude, the primary signal separating tissues in bulk RNA-seq, which aligns closely with the evaluation objective. Second, the evaluation samples overlap with its pretraining corpus, so performance partly reflects in-distribution familiarity. In contrast, TifBERT was pretrained without expression values or access to these samples and must rely on co-occurrence structure derived from TF-IDF ranked gene sequences. Because each disease corresponds to a unique tissue of origin, tissue prediction represents a coarser version of disease classification, explaining the similar trends across both metrics.

### Cross-Cancer Retrieval

Local embedding structure was evaluated using Precision@k, defined as the fraction of a sample’s k nearest neighbors (cosine distance) sharing the same disease label. BulkRNABert attained higher scores across all k. Because these samples were observed during pretraining, neighbor retrieval for BulkRNABert benefits from in-distribution similarity. Notably, the performance gap is smaller than in linear probing, suggesting that much of the linear probe advantage arises from globally aligned decision boundaries rather than substantially improved local biological consistency.

### Lung Subtype Discrimination

Discrimination between lung adenocarcinoma (n=26) and squamous cell carcinoma (n=23) was evaluated to isolate finer biological structure in models sharing tissue origin but differing histologically. A binary linear probe yielded comparable mean accuracy for both models, both exceeding the majority baseline (53.1%). However, BulkRNABert exhibited substantially larger variance across folds, indicating unstable subtype separation despite in-distribution exposure. The threefold reduction in variance for TifBERT (6.5% versus 19.2%) suggests that TifBERT representations provide more stable separation of histological subtypes despite the absence of in-distribution pretraining exposure, consistent with representations capturing regulatory structure beyond expression magnitude.

In summary, across evaluations, BulkRNABert performed strongly on tasks aligned with expression magnitude and benefited from in-distribution evaluation. Despite this advantage, TifBERT matched performance on intra-tissue subtype discrimination while exhibiting greater stability and substantially richer embedding geometry. The results indicated that the two models encoded distinct biological structure, with TifBERT producing representations that generalized better across datasets and normalization regimes, as supported by the GTEx transfer results.

## Discussion

The results presented here demonstrate that unordered high-dimensional expression profiles can be converted into informative gene sequences through sample-specific TF-IDF ranking, enabling transformer-based contextual learning without dependence on expression-value reconstruction. This finding has practical implications for bulk RNA-seq modeling: by decoupling pretraining from numerical expression scales, TifBERT produces representations that remain applicable across datasets generated with different normalization pipelines, a common challenge in multi-cohort transcriptomic studies. The consistent performance of a single pretrained encoder across cancer classification, pathway regression, expression prediction, and survival analysis further supports the feasibility of foundation-style modeling for bulk transcriptomics, a modality that remains central to clinical genomics but has received comparatively less attention in recent self-supervised representation learning.

A key finding is that TF-IDF ordering consistently outperformed raw-expression and Z-score-based ordering across multiple downstream tasks. This suggests that the IDF component provides information beyond expression magnitude by emphasizing genes that are selectively expressed across the cohort. In the context of bulk RNA-seq, where many highly expressed genes reflect housekeeping or broad tissue-level programs, this sample-specific specificity signal may help expose disease-relevant or pathway-relevant transcriptional structure to the encoder. The substantial gap between TifBERT and a non-contextual top-*K* TF-IDF baseline further indicates that performance is not explained by identifying highly ranked genes alone; masked gene modeling appears to learn contextual gene-to-gene relationships that improve downstream discrimination in a way that gene identity information without sequence context cannot.

In direct comparison with BulkRNABert, TifBERT achieved competitive subtype discrimination with substantially greater stability and markedly richer embedding geometry, despite operating without expression discretization, in-distribution pretraining exposure, or access to evaluation samples during training. This comparison suggests that the design choices introduced here, namely the avoidance of expression binning and the use of a sample-specific rather than globally fixed gene sequence, represent a meaningful advance over prior bulk transformer approaches. More broadly, TifBERT differs from approaches that depend on expression discretization, external gene embeddings, or landmark-gene restriction by learning directly from transcriptome-wide gene identities ordered in a sample-specific manner. This design reduces dependence on dataset-specific preprocessing choices and may improve portability across normalization schemes and cohorts, particularly in settings where expression scales differ substantially across studies.

Several limitations should be addressed in future work. First, although TCGA provides a large and diverse pan-cancer training resource, broader validation across independent disease cohorts, sequencing platforms, and clinical study designs will be needed to establish generalizability beyond the TCGA Pan-Cancer Atlas. Second, the current external evaluation on GTEx demonstrates preservation of tissue-level transcriptomic structure, but future work should include task-specific prediction benchmarks on independent healthy tissue cohorts to more rigorously quantify cross-dataset transferability. Third, while TF-IDF ordering provides an interpretable input construction strategy, further work is needed to characterize the biological meaning of learned attention patterns, gene neighborhoods, and masked gene predictions, which may reveal the transcriptional co-regulatory structures captured during pretraining. Fourth, the divergence in survival prediction performance between overall survival and cancer-specific endpoints warrants investigation; overall survival captures all-cause mortality and may depend on expression signals that differ from those relevant to disease-specific or progression-based outcomes, and future work should examine whether endpoint-specific pretraining objectives or input representations could improve prognostic modeling across all endpoints.

In summary, TifBERT provides a normalization-independent framework for applying self-supervised transformer learning to unordered high-dimensional bulk RNA-seq profiles. By combining sample-specific TF-IDF gene ordering with masked gene identity prediction, the framework bridges a methodological gap between transcriptomic foundation models and the continued practical importance of bulk RNA-seq in cancer genomics, biomarker discovery, and translational cohort analysis. The results suggest that the key design choices, sample-specific gene ordering and decoupled pretraining from expression magnitudes, are broadly applicable to bulk transcriptomic datasets and warrant evaluation in additional disease and cohort settings.

## Code and Data Availability

All code, pretrained model weights, and experimental configurations are publicly available at https://github.com/mohsenh17/TifBERT.

Bulk RNA-seq data were obtained from the TCGA Pan-Cancer (PANCAN) cohort via the UCSC Xena platform (https://xenabrowser.net/datapages/?cohort=TCGA%20Pan-Cancer%20(PANCAN)). Survival endpoints were obtained from the TCGA Pan-Cancer Clinical Data Resource (TCGA-CDR)^31^, providing OS, PFI, DSS, and DFI labels. Cancer type labels were derived from the primary disease phenotype file. Pathway activity targets were obtained from z-scored ssGSEA scores for 1,387 PARADIGM pathways, also available through the Pan-Cancer Atlas Hub.

External healthy tissue data from GTEx were accessed via the UCSC Toil RNA-seq Recompute compendium and processed using the same harmonization pipeline as TCGA data. All datasets are publicly accessible and were used in accordance with their respective data access policies.

## Author Contributions

S.M.H. implemented the TifBERT framework, conducted all experiments, performed data analysis, and wrote the original draft. D.S. conceptualised, supervised the project, acquired funding, provided critical review of the methodology, and revised the manuscript. All authors read and approved the final manuscript.

## Acknowledgements

This study was funded by New Frontiers in Research Fund-Exploration NFRFE-2024-00669.

